# EpiSort: Enumeration of cell types using targeted bisulfite sequencing

**DOI:** 10.1101/677211

**Authors:** Dvir Aran, Ron S. Dover, Karen E. Lundy, Michael D. Leipold, Ji Xuhuai, Shana L. McDevitt, Mark M. Davis, Atul J. Butte

## Abstract

The cellular composition of tumors is now recognized as an essential phenotype, with implications to diagnostic, progression and therapy efficacy. A tool for accurate profiling of the tumor microenvironment is lacking, as single-cell methods and computational approaches are not applicable or suffer from low accuracy. Here we present *EpiSort*, a novel strategy based on targeted bisulfite sequencing, which allows the accurate enumeration of 23 cell types and may be applicable to cancer studies.

## Main Text

Recent years have seen a tremendous advancement in technologies that enable the investigation of cellular heterogeneity and composition of mixed tissues. While newer methods have been developed for single-cell profiling after tissue dissection, innovative computational deconvolution techniques have also been created that enable digital dissection of bulk tissues. The need for such technologies is now indisputable, and a growing body of evidence shows the importance of the cellular heterogeneity in diagnosis, prognosis, treatment efficacy, and for the development of novel treatment strategies in an array of common and complex diseases.^1^ In the study of cancer, more and more evidence shows that the complex milieu of the tumor microenvironment is playing a major role in both promoting and inhibiting the tumor’s growth, to invasion and to metastasize.^2^ It is thus clear that accurate portrayal of the cellular composition of the tumor is an important phenotype is crucial for improving existing treatments, for the discovery of predictive biomarkers, and for the development of novel therapeutic strategies.

While cellular composition can be profiled using single-cell technologies such as flow cytometry, it is a far more challenging task in solid tissues. Dissociation of solid tissues leads to destruction of the niche environment, thereby affecting cellular state and integrity. It is thus unclear if the relative proportion of cell types is maintained following the dissociation step. Another disadvantage of single-cell methods is the need to perform the analysis on fresh tissues, which requires a supporting operational system and may not be suitable for use in clinical settings. The emerging use of single-cell RNA sequencing may offer novel insights regarding internal cellular states, but for profiling the cellular composition it does not provide a suitable solution.

On the other hand, computational deconvolution techniques employ genomic profiling of the bulk tissue, and thus, in theory, can more reliably detect the actual, non-harmed, tissue composition.^3^ However, the accuracy of these methods is questionable, specifically in tumors. Gene expression profiles are highly variable, and the large dynamic range is highly susceptible to batch effects and cell states changes that make it an impossible task to accurately predict the abundance of cell types.

Another genomic profiling measurement used for deconvolution is DNA methylation.^4^ DNA methylation, like gene expression, is cell type specific,^5^ however its scales are fractions and with linear association with the mixture’s composition, and not continuous as in gene expression, making it a more potent measurement for portraying the tissue composition. A number of methods have employed DNA methylation arrays to predict the composition of handful of immune subsets,^4,6,7^. However, only major immune subsets can be assessed since the limited breadth of methylation arrays is incapable of distinguishing closely related cell types. In addition, the accuracy of these methods is yet insufficient, predominantly due to the inability of individual CpGs to distinguish between stochastic and real bimodal epipolymorphism (Figure 1a).^8^ Accurate bisection of genomic locus to epistates is possible with bisulfite sequencing, and could therefore allow for reliable deconvolution of mixtures. Accordingly, whole genome bisulfite sequencing (WGBS) has been used for purity estimations of tumors,^9^ however, it cannot be used for identifying cell types with low abundance due to the low read depth and its high cost.

**Figure 1.**
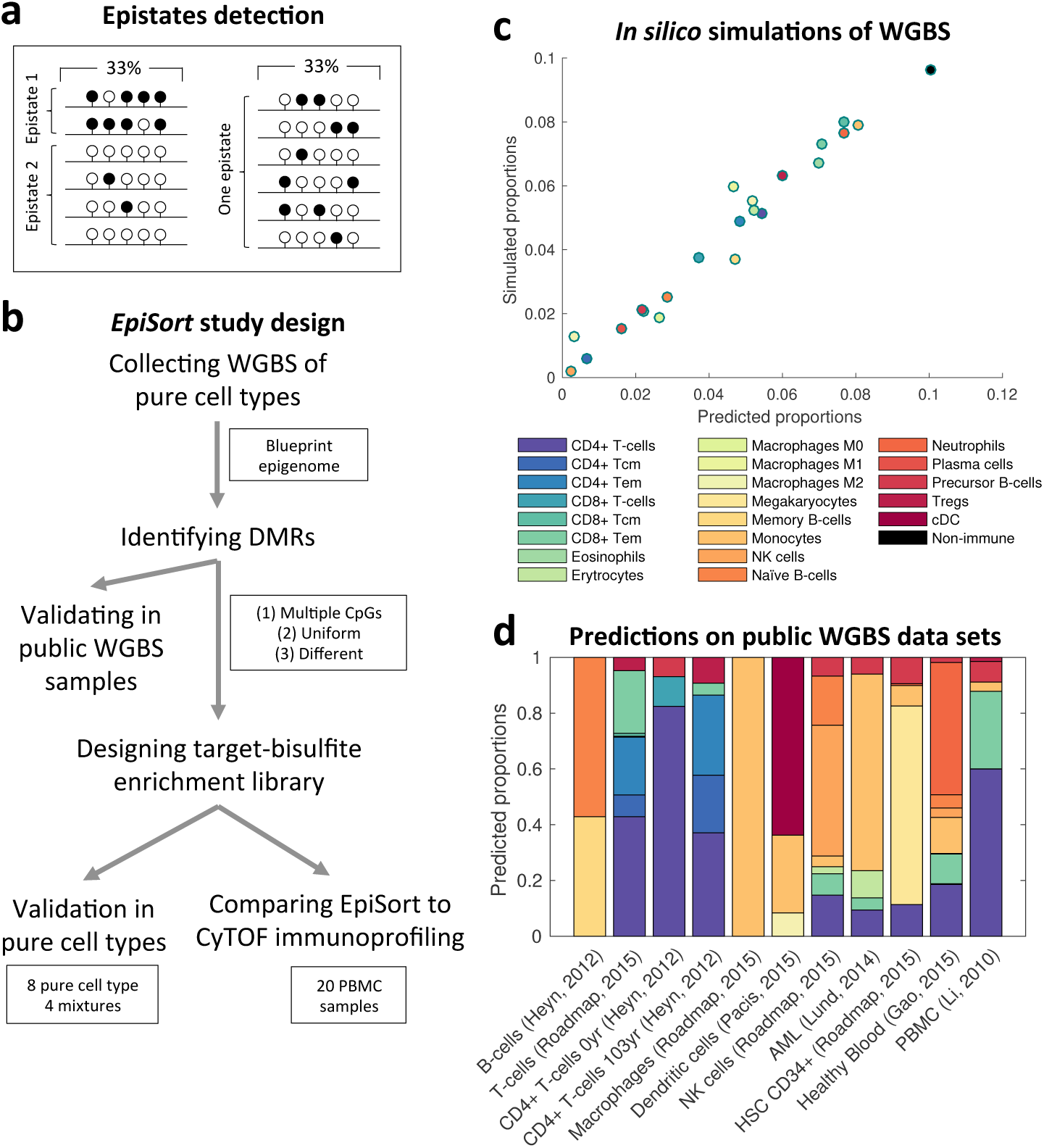
Designing a DNA methylation-targeting library for deconvolution of mixed tissues. **A**. A toy example of the problem of relying on single CpG to infer epistates. In both examples a third of the CpGs are methylated (black). In the left reads set it is clear that there are two epistates, possibly from two different cell types, while the right reads set cannot be split to epistates. However, a method that will call methylation from independent CpGs will not be able to differentiate between the two sets. **B**. An overview of the design and validation of EpiSort. **C**. An example of inferences of underlying abundance of an in-*silico* simulation based on the WGBS data. **D**. Inferring cell proportions in public WGBS samples. Colors are as in C. CD4+ T-cells of a centenarian are more specialized then newborn’s, possibly explaining many differences found in the methylation profiles. A mislabeled macrophages sample was detected as 100% monocytes.

Here we present *EpiSort*, a novel method to accurately enumerate 23 cell types in mixed tissues, using massive deep bisulfite sequencing of targeted loci specifically chosen to discriminate between cell types (Figure 1b). *EpiSort* targets cell types specific loci with multiple CpGs that allow a clear distinction of epistates. To identify these loci, we collected 60 WGBS samples, corresponding to 21 immune cell types from the Blueprint epignome project^10^ and two WGBS representing fibroblasts (IMR90)^11^ and epithelial cells (HMEC)^12^, totaling 23 cell types (Supplementary Table 1). Using the methylation fraction per CpG, we then scanned the genome to detect informative loci that will allow discriminating between cell types. Our search yielded 9,291 uniform differentially methylated regions (UDMRs) (Supplementary Table 2). We first tested our UDMRs capability to predict cell types composition in simulated *in silico* mixtures. We performed simulations by generating a pool of *in silico* bisulfite reads based on the underlying mixture of the cell types, and randomly choosing 200 reads from the pool per region. Our analyses showed highly accurate inferences of underlying abundances, differentiating closely related cell types, and the capacity to identify extremely low abundant cell types (Figure 1c).

We next applied our reference UDMR matrix on publicly available WGBS of pure and mixed samples obtained from MethBase.^13^ Accurate predictions are not expected in this type of analysis due to low read depth intrinsic in WGBS experiments, and also because the raw reads were not available, and thus calls were made based on individual CpGs in the locus. Despite this, we were able to detect the cell type in all cases, and detect known tendencies in peripheral blood mononuclear cells (PBMCs) (Figure 1d and Supplementary Table 3). Interestingly, one data set described as macrophages was called as 100% monocytes by our method. Tracking the source of this data revealed that these are indeed CD14+ monocytes (GSM1186661), and were misannotated in MethBase.

Next, we designed a custom DNA enrichment kit suitable for bisulfite sequencing to allow for targeting of the UDMRs. We applied the enrichment assay on DNA extracted from 8 pure immune cell types. The targeting assay yielded a low on-target rate, ranging in 10-15% of mapped reads, possibly due to using universal blocking oligos. Thus, per sample approximately only 300K reads were available (Supplementary Table 4), and only approximately 10% of the targeted loci were sequenced in all samples at least 20 times. Yet, the analysis of the pure cell types showed that the majority of UDMRs are concordant with the Blueprint-reference matrix, are uniform and have low uninformative rate (Supplementary Figure 1). However, the concordance of unmethylated regions in neutrophils and plasmacytoid dendritic cells (pDCs) was low, indicating that enumeration of these cell types may not be accurate. The dendritic cells in Blueprint are from conventional DC, which may explain the incompatibility we observe with pDC.

We next employed the 8 pure immune cell types to perform simulated *in silico* mixtures, and predicted the underlying abundances using the original reference matrix learned from the Blueprint dataset. These simulations are based on the authentic sequenced reads and not on individual CpGs as in previous simulations, and therefore methylation calls may be partial and uninformative, making these simulations resembling real data. The analysis revealed that in all cell types, excluding pDCs, the predictions were highly accurate (Figure 2a). We also generated 4 synthetic *in vitro* mixtures from the 8 cell types, with varying proportions of the cell types. In all 4 mixtures, *EpiSort* was able to recover significantly the underlying abundances (p *value* < 0.05) (Figure 2a).

**Figure 2.**
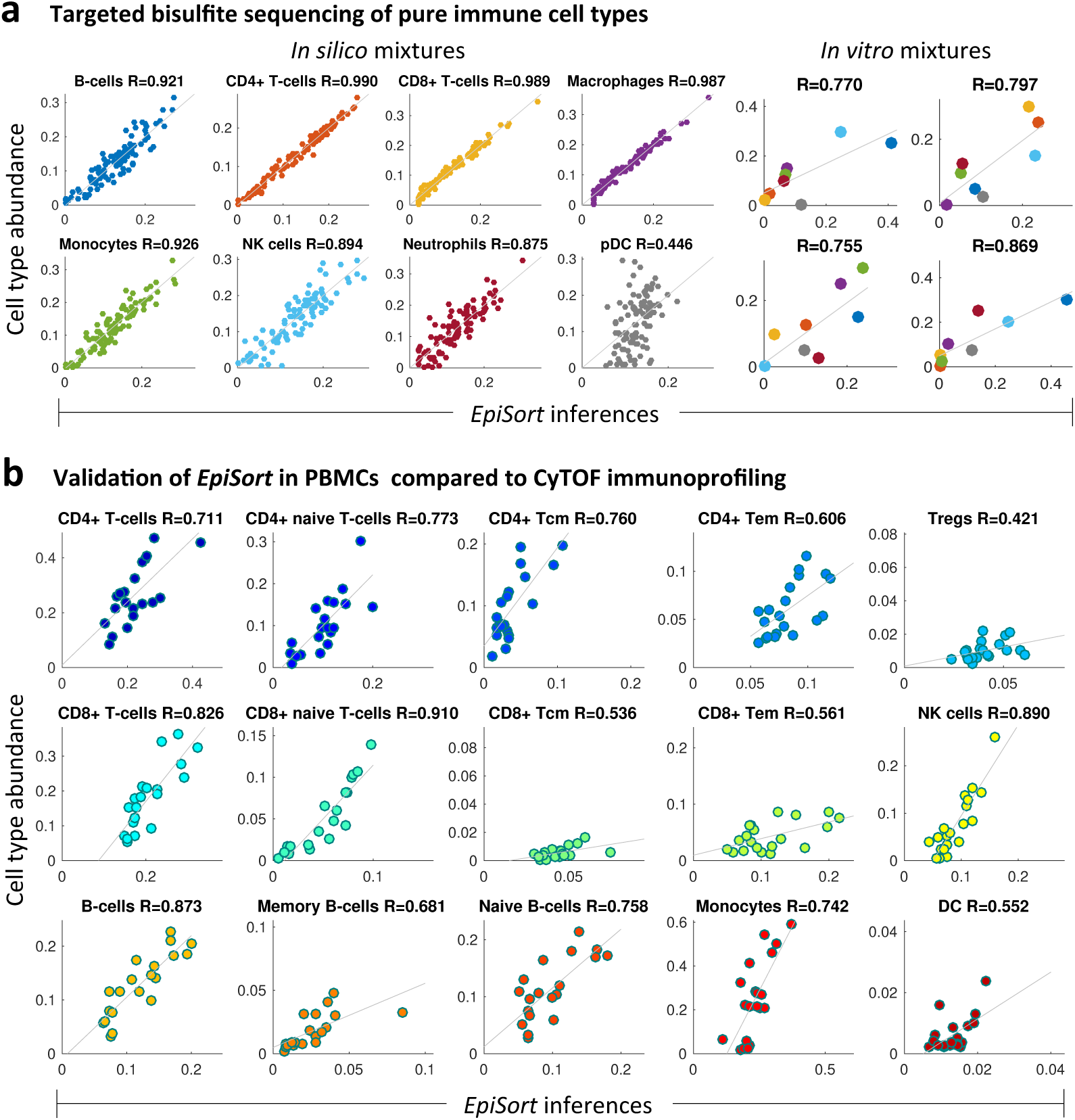
Evaluation of EpiSort predictive power. **A**. Applying *EpiSort* on 8 pure immune cell types. **Left:** Inferences from 100 in-*silico* mixtures of the cell types. The reference dendritic cell is convenctional (cDC), while we applied it on plasmacytoid (pDC), presumably explaining its prediction failure. A linear function was used to shift and scale the associations. **Right:** Inferences of the cell types abundances in 4 in-vitro mixtures of the 8 cell types. The inferences were shifted and scaled according to the parameters learned from the simulations. Colors of data points correspond to colors of cell types in A. **B**. Applying *EpiSort* on 20 PBMCs, and comparing the inferences with abundances measured using CyTOF immunoprofiling on the same samples. In 14 of the 15 cell types the correlation is significant (p-*value* < 0.05), and in regulatory T-cells (Tregs) the significance is marginal (p=0.06).

Finally, we test *EpiSort* against the gold-standard benchmark of cell counting. We performed the *EpiSort* assay on the DNA extracted from 20 PBMC samples, which were also analyzed by mass spectrometry (CyTOF) immunoprofiling. The DNA for this experiment was extracted from TRIzol samples and was of low quality; for some of the samples very low amount of DNA was available (Supplementary Table 4). Yet, we found that the signal from this experiment was sufficient to infer with high accuracy most of the cell types in the mixture (Figure 2b). Lower reliability was only observed in low abundant cell types, presumably related to non-sufficient on-target reads. Finally, we compared the correlations of our inferences with gene expression-based methods, including CIBERSORT^14^ and our own xCell,^15^ and found that *EpiSort* is more reliable in inferring the proportions of cell types in PBMCs (Supplementary Figure 2).

We believe that *EpiSort* can serve as a potent tool for tissue composition analysis, and in particular for studying tumor microenvironment heterogeneity. The advantage of *EpiSort* over single-cell based technologies is clear: since no cell suspension is required, the analysis could be performed on solid tissues without additional destructive dissociation steps, and importantly, it could be performed on archived samples. In cancer studies, this is crucial. DNA methylation analysis from formalin-fixed paraffin-embedded (FFPE) tissues is possible,^16^ and thus *EpiSort* provides a compelling tool for reexamining archived cohorts.

We have shown here that *EpiSort* is more accurate than other bulk tissue-based methods. We expect that the accuracy can be further significantly improved with better optimization of the on-target reads. In addition, the reference matrix can be optimized as well by analyzing pure cell types with *EpiSort*, and removing inaccurate loci. However, since *EpiSort* relies on a reference matrix it is constrained by our current understanding of cell types annotations, and cannot detect continuums. It should be emphasized that *EpiSort* is only useful for enumerating cell types and not for any purpose beyond that.

In summary, *EpiSort* is an accurate low cost method to enumerate cell populations in a bulk mixture. It can be performed with low quality and low amount of input DNA, and with high accuracy compared to other methods. We prospect that *EpiSort* will be a valuable tool for identifying predictive biomarkers that could lead to personalized cancer immunotherapy strategies, and facilitate new cancer immunotherapy approaches.

## Acknowledgments

This work was supported by the Gruss Lipper Postdoctoral Fellowship to D.A., and the National Cancer Institute (U24 CA195858) and the National Institute of Allergy and Infectious Diseases (Bioinformatics Support Contract HHSN272201200028C) to A.J.B. The content is solely the responsibility of the authors and does not necessarily represent the official views of the National Institutes of Health.

## Methods

### Identifying uniform differentially methylated regions

DNA methylation calls of human whole genome bisulfite sequencing samples mapped to hg38 were downloaded from the Blueprint Epigenome project^10^ and from MethBase.^13^ Altogether we obtained 64 samples, which were aggregated to 23 cell types. Two samples were removed due to low coverage. We scan the genome for *N* consecutive CpGs in a region <100bp, such that in each cell type all CpGs are methylated or unmethylated. We measure this characteristic, which we term uniformity, using the following formula:

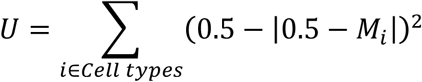

Then for each pair of cell types we use calculate the difference (*D*) of each of the CpGs in the region. We use different threshold for *N, U* and *D*, starting from stringent thresholds and gradually relaxing the thresholds to allow 100 regions for each per of cell types. Ranges are N=5 to 2, U=0.01 to 0.05 and D=0.95 to 0.5. Regions containing CpGs that are unavailable in more than 2 cell types are removed. The search yielded 9,291 regions across the genome.

### CyTOF Immunoprofiling

PBMCs of healthy volunteer were collected by the Stanford Blood Center, and were provided with courtesy from the Institute for Immunity, Transplantation and Infection (HIMC) at Stanford University.. PBMCs were analyzed using the CyTOF immunoprofiling protocol described in Leipold & Maeker.^17^ Normalization of the mass cytometry data was performed according to Finck et al.^18^ Next, normalization beads, debris, doublets, and dead cells were removed from the data before marker-based gating to determine the cell populations. The fractions presented are the number of cells in a particular gated subpopulation divided by the number of viable singlet cells. The remainders of the donor cells not used for mass cytometry analysis were preserved with TRIzol and DNA was extracted using…

### DNA from pure immune cell types in peripheral blood

White blood cell concentrates of TrimaAccel® leukocyte reduction system (LRS) cones recovered after Plateletpheresis procedure were obtained from two donors by the Stanford Blood Center according to Institutional Review Board (IRB) (“Minimal Risk Research Related Activities at Stanford Blood Center”, Protocol #13942) protocol. Pure cell populations were isolated using EasySep Human negative selection kits (STEM CELL technologies, >95% purity). In addition, we analyzed genomic DNA from frozen human peripheral blood Macrophages (≥ 90%) and Plasmacytoid Dendritic Cells (pDCs, (≥ 85%)). DNA was purified using QIAamp DNA Mini Kit (QIAGEN).

### Targeted bisulfite-sequencing

Extracted genomic DNA (gDNA) was sent to the University of California-Berkeley, Institute for Quantitative Biosciences (QB3), Functional Genomics Laboratory where it was prepared for Illumina short-read sequencing using a custom designed Nimblegen targeted capture probe panel. The DNAs were initially spiked with bisulfite conversion control, as indicated in the Nimblegen SeqCap Epi enrichment protocol (Roche Diagnostics Corp.,Indianapolis, IN) and sheared to 200 base-pairs (bp) on a Covaris S220 Focused-Ultrasonicator (Covaris, Woburn, MA). Fragmented DNA was cleaned and concentrated with MinElute® PCR Purification columns (QIAGEN Inc.,Valencia, CA) and prepared for sequencing on an Apollo 324™ with PrepX™ ILM 32i DNA Library Kits (WaferGen Biosystems, Fremont, CA), using eight base-pair dual-indexed methylated adapters (Integrated DNA Technologies Inc., Chicago, IL). Post library preparation, bisulfite conversion was performed on the adapter-ligated, indexed libraries using the EZ DNA Methylation-Lightning kit (Zymo Research Corp., Irvine, CA), after which bisulfite-converted libraries were amplified with 12 polymerase chain reaction (PCR) cycles using KAPA HiFi Hotstart Uracil+ ReadyMix and Ligation-Mediated PCR oligos (Roche Diagnostics Corp.,Indianapolis, IN). Samples submitted with less than one microgram of extracted gDNA were amplified with an additional 2 PCR cycles to increase concentration pre-capture. Amplified libraries were purified using AMpure XP solid-phase reversible immobilization beads (Beckman Coulter Inc., Indianapolis, IN) and quantified using a high sensitivity DNA Qubit assay (Thermo Fisher Scientific, Billerica, MA).

Eight one-microgram pools, consisting of four different 250ng aliquots of purified, bisulfite-converted library, were hybridized to capture probes for approximately 70 hours following the SeqCap Epi Enrichment system protocol (Roche Diagnostics Corp.,Indianapolis, IN). Bisulfite capture enhancer provided in the capture kit and xGen universal oligo blockers (Integrated DNA Technologies Inc., Chicago, IL) were added to the pools prior to probe hybridization. Capture bead-bound sample pools were washed and amplified for 16 PCR cycles using KAPA HiFi Hostart ReadyMix and LM-PCR oligos (Roche Diagnostics Corp.,Indianapolis, IN). Final products were visualized on a high-sensitivity DNA Advanced Analytical Fragment Analyzer assay (Advanced Analytical Technologies Inc., Ames, IA). Pools were then transferred to the QB3 Genomics Sequencing Laboratory at UC Berkeley and quantified in duplicate on a BioRad CFX connect quantitative PCR thermal cycler (BIO-RAD Laboratories, Hercules, CA) using the KAPA Universal qPCR Mix Library Quant Kit for Illumina (Roche Diagnostics Corp.,Indianapolis, IN). The eight capture pools were then pooled equimolar and sequenced across two Illumina HiSeq2500 v2 150bp single-end rapid lanes with a ten percent PhiX control library spike (Illumina Inc., San Diego, CA). Basecall files (bclfiles) were demultiplexed and converted to fastQ file format using the bcl2fastq v2.18 software (Illumina Inc., San Diego, CA).

### EpiSort pipeline

#### Reference matrix

Based on the WGBS datasets we generated a 9,291×23 binary matrix for each of the UDMRs and cell types. Each cell in the matrix is 1 if the regions methylation > 0.5, and 0 otherwise. The reference matrix can be further refined using targeted bisulfite sequencing of pure cell types, which will yield higher reliability due to superior read depth.

#### DNA extraction

The requirement for the targeting assay is 1 microgram of DNA. However, our analyses show that the targeting assay can be performed with very low amount of DNA, as we were able to recover the composition of a sample were only 40 nanograms were available.

#### Targeting

We used here the SeqCap Epi kit to target 9,291 regions, multiplexing 4 samples per reaction.

#### Sequencing

All 32 samples were multiplexed and sequenced together using one HiSeq 2500 lane. We used 150bp single-read sequences.

#### Alignment

Bismark Bisulfite Mapper^19^ was used to align the reads to the hg38 reference genome. Using Matlab sparse matrices we efficiently analyzed each sample to count the number of methylated, unmethylated and uninformative reads in the targeted regions. A read is called methylated if the majority of its CpGs are methylation, unmethylated if the majority is unmethylated, and uninformative if equal number of methylated and methylated CpGs.

#### Analysis

For the input sample we calculate for each region the methylation call, which is the fraction of methylated reads from all informative reads. This generates a vector (*b*), and with the reference matrix (*A*) we perform linear regression analysis to compute the underlying abundances of the cell types (*x*). To perform the analysis we employ robust fitting to reduce the effect of outliers.^20^

## References

1. Klemm, F. & Joyce, J. A. Microenvironmental regulation of therapeutic response in cancer. Trends Cell Biol. 25, 198–213 (2015).

2. Hanahan, D. & Coussens, L. M. Accessories to the Crime: Functions of Cells Recruited to the Tumor Microenvironment. Cancer Cell 21, 309–322 (2012).

3. Newman, A. M. & Alizadeh, A. A. High-throughput genomic profiling of tumor-infiltrating leukocytes. Curr. Opin. Immunol. 41, 77–84 (2016).

4. Accomando, W. P., Wiencke, J. K., Houseman, E. A., Nelson, H. H. & Kelsey, K. T. Quantitative reconstruction of leukocyte subsets using DNA methylation. Genome Biol 15, R50 (2014).

5. Baron, U. et al. DNA methylation analysis as a tool for cell typing. Epigenetics 1, 55–60 (2006).

6. Houseman, E. A. et al. DNA methylation arrays as surrogate measures of cell mixture distribution. BMC Bioinformatics 13, 86 (2012).

7. Heiss, J. A. et al. Training a model for estimating leukocyte composition using whole-blood DNA methylation and cell counts as reference. Epigenomics 9, 13–20 (2017).

8. Landan, G. et al. Epigenetic polymorphism and the stochastic formation of differentially methylated regions in normal and cancerous tissues. Nat. Genet. 44, 1207–14 (2012).

9. Zheng, X. et al. MethylPurify: tumor purity deconvolution and differential methylation detection from single tumor DNA methylomes. Genome Biol. 15, 419 (2014).

10. Blueprint Epigenome project: http://www.blueprint-epigenome.eu/

11. Lister, R. et al. Human DNA methylomes at base resolution show widespread epigenomic differences. Nature 462, 315–322 (2009).

12. Hon, G. C. et al. Global DNA hypomethylation coupled to repressive chromatin domain formation and gene silencing in breast cancer. Genome Res. 22, 246–258 (2012).

13. Song, Q. et al. A Reference Methylome Database and Analysis Pipeline to Facilitate Integrative and Comparative Epigenomics. PLoS One 8, e81148 (2013).

14. Newman, A. M. et al. Robust enumeration of cell subsets from tissue expression profiles. Nat. Methods 12, 453–7 (2015).

15. Aran, D., Hu, Z. & Butte, A. J. xCell: Digitally portraying the tissue cellular heterogeneity landscape. bioRxiv 114165 (2017). doi:10.1101/114165

16. Dumenil, T. D. et al. Genome-wide DNA methylation analysis of formalin-fixed paraffin embedded colorectal cancer tissue. Genes. Chromosomes Cancer 53, 537–548 (2014).

17. Leipold, M. & Maecker, H. Phenotyping of Live Human PBMC using CyTOFTM Mass Cytometry. BIO-PROTOCOL 5, (2015).

18. Finck, R. et al. Normalization of mass cytometry data with bead standards. Cytom. Part A 83A, 483–494 (2013).

19. Krueger, F. & Andrews, S. R. Bismark: a flexible aligner and methylation caller for Bisulfite-Seq applications. Bioinformatics 27, 1571–2 (2011).

20. Buja, A., Tukey, P. & University of Minnesota. Institute of Mathematics and its Applications. Computing and graphics in statistics. Computing and graphics in statistics (Springer-Verlag, 1991). at <http://dl.acm.org/citation.cfm?id=140809>

