## Supplementary Figures and Tables for "EpiSort: Enumeration of cell types using targeted bisulfite sequencing"

### Supplementary Table 1:
#### Whole genome bisulfite sequencing samples used to identify uniform differentially methylated regions

| Cell types | # of WGBS samples |
|---|---|
| **CD4+ T-cells** | 3 |
| **CD4+ Tcm** | 3 |
| **CD4+ Tcm** | 3 |
| **CD4+ Tem** | 1 |
| **CD8+ T-cells** | 4 |
| **CD8+ Tcm** | 1 |
| **cDC** | 2 |
| **Eosinophils** | 1 |
| **Erytrocytes** | 2 |
| **Macrophages M0** | 7 |
| **Macrophages M1** | 6 |
| **Macrophages M2** | 5 |
| **Megakaryocytes** | 1 |
| **Memory B-cells** | 1 |
| **Monocytes** | 6 |
| **Naïve B-cells** | 5 |
| **Neutrophils** | 6 |
| **NK cells** | 2 |
| **Plasma cells** | 2 |
| **Precursor B-cells** | 1 |
| **Tregs** | 1 |
| **Fibroblasts (IMR90)** | 1 |
| **Endothelia cells (HMEC)** | 1 |

**Table S1. Whole genome bisulfite sequencing samples used to identify uniform differentially methylated regions.** Methylation calls of immune cell types was downloaded from the Blueprint Epigenome Project [1], IMR90 from [2] and HMEC from [3].

### Supplementary Table 3:

#### Deconvolving publicly available whole genome bisulfite sequencing data

| | B-cells (Heyn, 2012) | T-cells (Roadmap, 2015) | New born CD4+ T-cells (Heyn, 2012) | Ceneterian CD4+ T-cells (Heyn, 2012) | Macrophages (Roadmap, 2015) | Dendritic cells (Pacis, 2015) | NK cells (Roadmap, 2015) | AML (Lund, 2014) | HSC CD34+ (Roadmap, 2015) | Healthy Blood (Gao, 2015) | PBMC (Li, 2010) |
|---|---|---|---|---|---|---|---|---|---|---|---|
| **CD4+ T-cells** | 0 | 0.4288 | 0.8245 | 0.371 | 0 | 0 | 0.1473 | 0.0943 | 0.1139 | 0.1859 | 0.5997 |
| **CD4+ Tcm** | 0 | 0.0779 | 0 | 0.2066 | 0 | 0 | 0 | 0 | 0 | 0 | 0 |
| **CD4+ Tem** | 0 | 0.2073 | 0 | 0.2869 | 0 | 0 | 0 | 0 | 0 | 0 | 0.0008 |
| **CD8+ T-cells** | 0 | 0.0031 | 0.1063 | 0 | 0 | 0 | 0 | 0 | 0 | 0.0026 | 0 |
| **CD8+ Tcm** | 0 | 0.0105 | 0 | 0 | 0 | 0 | 0 | 0 | 0 | 0 | 0 |
| **CD8+ Tem** | 0 | 0.2251 | 0 | 0.0429 | 0 | 0 | 0.077 | 0.0435 | 0 | 0.1067 | 0.2776 |
| **Eosinophils** | 0 | 0 | 0 | 0 | 0 | 0 | 0 | 0 | 0 | 0 | 0 |
| **Erytrocytes** | 0 | 0 | 0 | 0 | 0 | 0 | 0.025 | 0.097 | 0 | 0.002 | 0 |
| **Macrophages M0** | 0 | 0 | 0 | 0 | 0 | 0 | 0 | 0 | 0 | 0 | 0 |
| **Macrophages M1** | 0 | 0 | 0 | 0 | 0 | 0 | 0 | 0 | 0 | 0 | 0 |
| **Macrophages M2** | 0 | 0 | 0 | 0 | 0 | 0.0838 | 0 | 0 | 0 | 0 | 0 |
| **Megakaryocytes** | 0 | 0 | 0 | 0 | 0 | 0 | 0 | 0 | 0.7119 | 0 | 0 |
| **Memory B-cells** | 0.4289 | 0 | 0 | 0 | 0 | 0 | 0 | 0 | 0 | 0 | 0 |
| **Monocytes** | 0 | 0 | 0 | 0 | 1 | 0.2786 | 0.0386 | 0.7049 | 0.0736 | 0.1287 | 0.0332 |
| **NK cells** | 0 | 0 | 0 | 0 | 0 | 0 | 0.4692 | 0 | 0 | 0.034 | 0 |
| **Naive B-cells** | 0.5711 | 0 | 0 | 0 | 0 | 0 | 0.1758 | 0 | 0 | 0.0473 | 0 |
| **Neutrophils** | 0 | 0 | 0 | 0 | 0 | 0 | 0 | 0 | 0.006 | 0.4741 | 0 |
| **Plasma cells** | 0 | 0 | 0 | 0 | 0 | 0 | 0 | 0 | 0 | 0 | 0 |
| **Precursor B-cells** | 0 | 0 | 0.0692 | 0 | 0 | 0 | 0.0671 | 0.0602 | 0.0947 | 0.0187 | 0.0743 |
| **Tregs** | 0 | 0.0474 | 0 | 0.0927 | 0 | 0 | 0 | 0 | 0 | 0 | 0.0144 |
| **cDC** | 0 | 0 | 0 | 0 | 0 | 0.6376 | 0 | 0 | 0 | 0 | 0 |

**Table S3. Deconvolving publicly available whole genome bisulfite sequencing data.** Methylation calls were downloaded from MethBase [4]. The data was generated by [5,6,7,8,9,10]. Average methylation of the 9,291 regions was calculated for each sample (were data was available), and linear least square analysis was performed using the reference matrix we generated.

**B-cells (Heyn, 2012)** – normal B cells of an healthy donor (female, 4 y) were immortalized by Epstein-Barr-Virus (EBV), which explains the high number of memory B-cells.

**T-cells (Roadmap, 2015)** - all inferences are of T-cells, more CD4+ than CD8+, as expected.

**Newborn CD4+ T-cells (Heyn, 2012)** – majority of inferences from CD4+ naïve T-cells.

**Ceneterian CD4+ T-cells (Heyn, 2012)** – from over 80% of naïve CD4+ T-cells in the newborn to less than 40%, and now a majority of memory and regulatory T-cells. This specialization is expected [ref].

**Macrophages (Roadmap, 2015)** – our method detected 100% monocytes. This sample is labeled as macrophages in MethBase, however Roadmap protocols clearly states it is CD14+ monocytes.

**Dendritic cells (Pacis, 2015)** – majority was detected as dendritic cells.

**NK cells (Roadmap, 2015)** – not pure detection as in other samples, yet, only cell type that is detected is NK cells.

**AML (Lund, 2014)** – as expected, the majority of this myeloid leukemia sample is monocytes.

**HSC CD34+ (Roadmap, 2015)** – while the Blueprint project sequenced only megakaryocytes, we wondered wether our method will detect hematopoietic stem cells. Indeed, the majority of the sample was detected as megakaryocytes.

**Healthy Blood (Gao, 2015)** - blood is known to contain ~50% of neutrophils, ~30% T-cells (twice more CD4+ than CD8+), ~10% monocytes, 5% B-cells and ~3 NK cells [ref]. Our inferences are very close to this premise.

**PBMC (Li, 2010)** – PBMCs do not contain neutrophils, as our method detected.

### Supplementary Table 4:
#### Targeted bisulfite sequencing experiment

| Sample ID | nanograms of DNA | # of reads | # of mapped reads | # of on-target reads | # of on-target reads > 1 CpGs | % of mapped reads | % of on-target reads | % of on-target CpG>1 |
|---|---|---|---|---|---|---|---|---|
| B-cells | 1000 | 3,297,718 | 2,433,286 | 297,314 | 199,576 | 73.79% | 12.22% | 8.20% |
| CD4+ T-cells | 1000 | 7,692,226 | 5,494,052 | 966,632 | 662,952 | 71.42% | 17.59% | 12.07% |
| CD8+ T-cells | 1000 | 4,164,251 | 3,038,825 | 401,630 | 275,561 | 72.97% | 13.22% | 9.07% |
| Macrophages | 1000 | 4,888,903 | 3,533,893 | 373,949 | 251,605 | 72.28% | 10.58% | 7.12% |
| Monocytes | 1000 | 6,071,579 | 4,575,100 | 514,473 | 344,689 | 75.35% | 11.25% | 7.53% |
| Neutrophils | 1000 | 2,768,320 | 2,067,539 | 455,520 | 308,712 | 74.69% | 22.03% | 14.93% |
| NK cells | 1000 | 5,769,113 | 4,300,726 | 239,941 | 160,902 | 74.55% | 5.58% | 3.74% |
| pDCs | 1000 | 4,707,570 | 3,467,397 | 383,360 | 260,587 | 73.66% | 11.06% | 7.52% |
| Mix1 | 1000 | 5,573,496 | 4,218,053 | 448,145 | 299,564 | 75.68% | 10.62% | 7.10% |
| Mix2 | 1000 | 5,635,386 | 4,245,298 | 451,050 | 302,922 | 75.33% | 10.62% | 7.14% |
| Mix3 | 1000 | 5,475,612 | 3,922,760 | 460,612 | 312,168 | 71.64% | 11.74% | 7.96% |
| Mix4 | 1000 | 5,998,694 | 4,446,846 | 480,998 | 323,507 | 74.13% | 10.82% | 7.27% |
| CyTOF_1 | 1000 | 6,610,183 | 2,780,484 | 287,505 | 190,902 | 42.06% | 10.34% | 6.87% |
| CYTOF_2 | 1000 | 4,910,432 | 3,531,369 | 693,019 | 475,602 | 71.92% | 19.62% | 13.47% |
| CYTOF_3 | 1000 | 6,570,582 | 3,588,530 | 571,396 | 389,234 | 54.62% | 15.92% | 10.85% |
| CYTOF_4 | 364 | 5,161,766 | 1,978,904 | 429,171 | 297,452 | 38.34% | 21.69% | 15.03% |
| CYTOF_5 | 1000 | 6,513,762 | 2,558,167 | 575,027 | 399,591 | 39.27% | 22.48% | 15.62% |
| CYTOF_6 | 1000 | 5,198,847 | 2,048,256 | 438,047 | 300,801 | 39.40% | 21.39% | 14.69% |
| CYTOF_7 | 1000 | 9,250,237 | 3,391,783 | 616,873 | 420,119 | 36.67% | 18.19% | 12.39% |
| CYTOF_8 | 944 | 10,883,118 | 4,326,056 | 970,550 | 674,181 | 39.75% | 22.43% | 15.58% |
| CYTOF_9 | 111 | 3,362,894 | 2,138,966 | 323,281 | 211,896 | 63.60% | 15.11% | 9.91% |
| CYTOF_10 | 726 | 10,126,171 | 4,418,367 | 932,034 | 643,528 | 43.63% | 21.09% | 14.56% |
| CYTOF_11 | 149.2 | 3,560,551 | 1,782,363 | 341,066 | 221,943 | 50.06% | 19.14% | 12.45% |
| CYTOF_12 | 348 | 2,945,804 | 1,921,413 | 240,841 | 156,679 | 65.23% | 12.53% | 8.15% |
| CYTOF_13 | 1000 | 14,226,000 | 7,297,669 | 1,194,655 | 819,460 | 51.30% | 16.37% | 11.23% |
| CYTOF_14 | 1000 | 9,610,522 | 5,196,946 | 782,494 | 534,489 | 54.08% | 15.06% | 10.28% |
| CYTOF_15 | 798 | 4,261,885 | 1,954,090 | 332,265 | 218,378 | 45.85% | 17.00% | 11.18% |
| CYTOF_16 | 188.4 | 13,490,343 | 4,610,063 | 755,257 | 528,064 | 34.17% | 16.38% | 11.45% |
| CYTOF_17 | 42.6 | 11,409,093 | 4,886,962 | 687,239 | 469,870 | 42.83% | 14.06% | 9.61% |
| CYTOF_18 | 956 | 10,259,503 | 5,950,351 | 766,443 | 522,686 | 58.00% | 12.88% | 8.78% |
| CYTOF_19 | 674 | 9,533,693 | 5,493,328 | 597,229 | 408,043 | 57.62% | 10.87% | 7.43% |
| CYTOF_20 | 334 | 7,865,311 | 3,583,734 | 465,724 | 316,792 | 45.56% | 13.00% | 8.84% |

**Table S4. Targeted bisulfite sequencing experiment.** We performed the targeted bisulfite experiment on 32 samples: 8 pure cell types, 4 mixes of those cell types and 20 PBMCs which were also analyzed using CyTOF immunoprofiling. All captured pools were sequenced using one HiSeq 2000 single-end rapid lane. The table shows the amount of DNA per sample, number of reads, mapped reads using Bismark, on-target reads with at least 1 CpG intersecting targeted reads, and with at least 2 CpGs. Samples with as low as 42 nanograms of initial DNA performed equally as samples with 1ug initial DNA. The targeting was lower than expected, possibly due to using unverisal blocking oligos.

**Supplementary Figure 1:**
Targeted bisulfite sequencing of pure immune cell types

**A** Correlation between reference methylation levels and targeted methylation calls

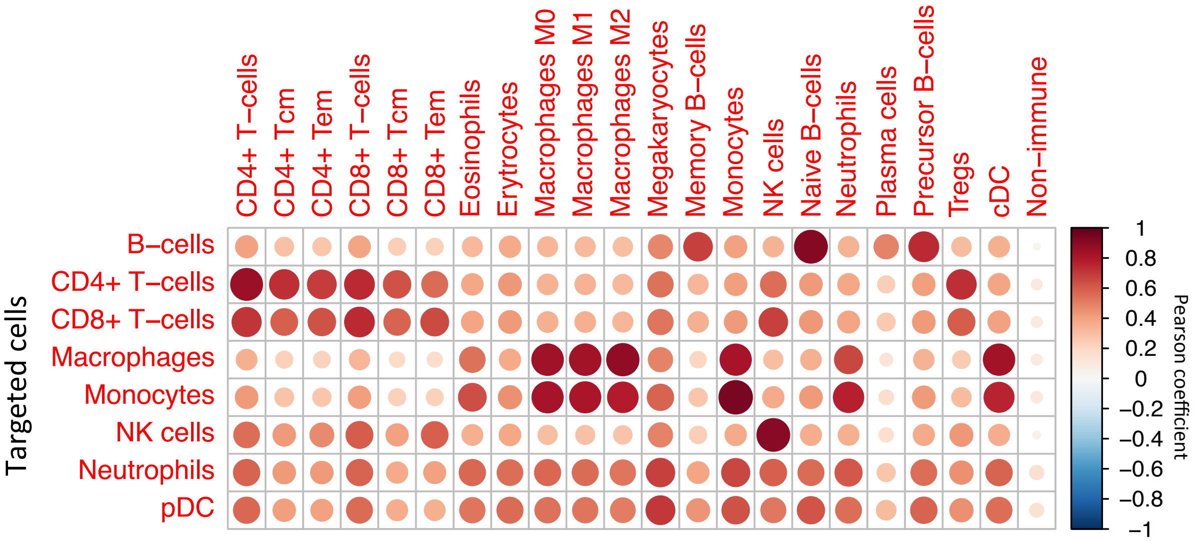


**B** Distribution of methylation calls of targeted cells

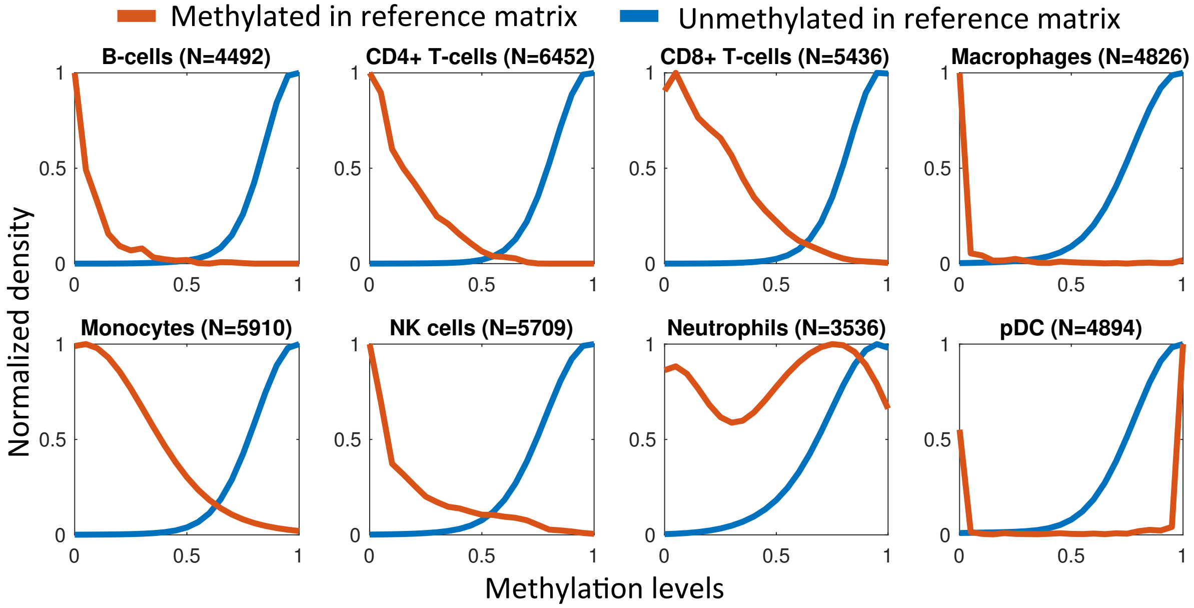

**C** Discordance between reference and targeting

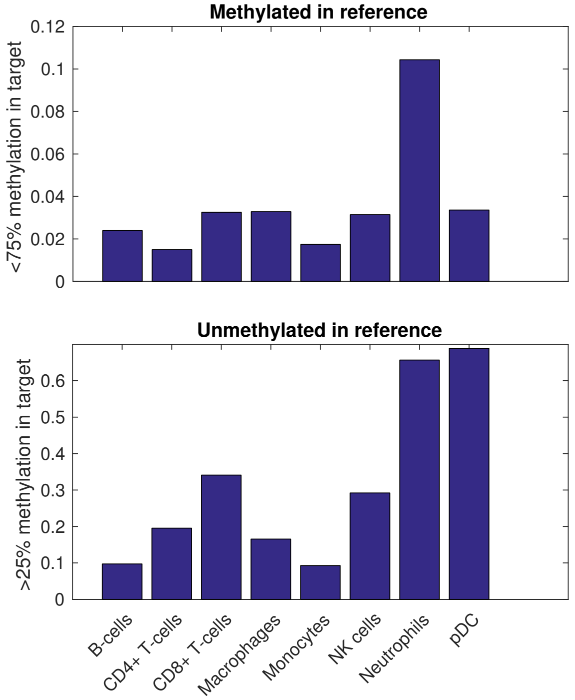


**D** Deconvolving the targeted samples

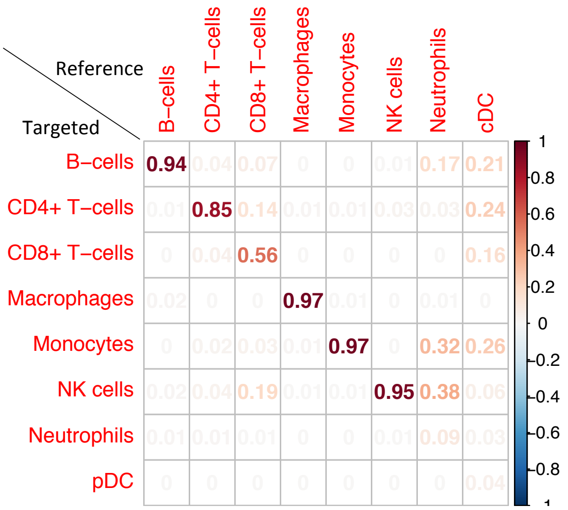


**Figure S1. Targeted bisulfite sequencing of pure immune cell types.** We performed targeted bisulfite sequencing on eight pure immune cell types. The percent of on-target reads was lower than expected, thus we were not able to use the full power of the methods. Only ~55% of the regions had at least 20 reads, and only 22% over 50 reads. We computed methylation calls for each of the regions, which is the fraction of reads with majority of CpGs methylated from all reads that have a majority. **A.** Pearson coefficients of correlations between the targeted methylation calls and the reference matrix. The top correlation for each of the targeted cells is the corresponding cell types. **B.** Using the 20 reads threshold, we plotted methylation calls for each of the regions (number of regions passing the threshold is shown for each cell type). The curves were split to those that were designated as methylated (blue) and unmethylated (red) in the reference matrix for the corresponding cell type. The reference for pDCs is cDCs, which might explain the failure of the unmethylated regions. **C.** Percent of discordance between the reference matrix and the targeted cells. **Top**: Percent of regions with methylation <75% in targeted cells from all methylated site in the reference matrix. **Bottom**: Percent of regions with methylation >25% in targeted cells from all unmethylated site in the reference matrix. High discordance in neutrophils and DCs. **D.** Inferences by EpiSort on the pure cell types. The predictions fails in both Neutrophils and pDCs, and also lower success in CD8+ T-cells.

### **Supplementary Figure 2:**
#### Comparison of EpiSort to gene-expression based methods

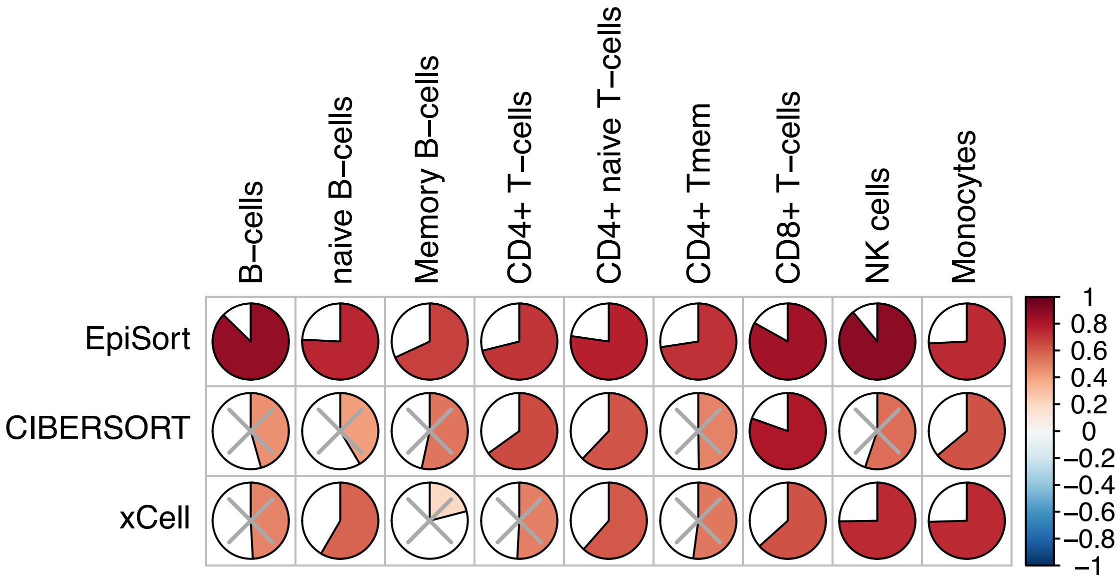


**Figure S2. Comparison of EpiSort to gene-expression based methods.** We did not have gene expression data on the same sample we performed the EpiSort analysis, thus we compared the correlations we obtained on 20the  samples we collected to inferences made on 20 samples collected by Newman et al. [11] (available in GSE65133). We reanalyzed the data with both CIBERSORT and xCell [12]. In all cell types comparison was possible, our inferences outperformed the gene-expression based inferences. Gray 'X' represent insignificant correlation (p-*value* > 0.01).

### Supplementary References


1. Blueprint Epigenome - Welcome to the Blueprint project at http://www.blueprint-epigenome.eu/
2. Lister, R. *et al.* Human DNA methylomes at base resolution show widespread epigenomic differences. *Nature* **462,** 315–322 (2009).
3. Hon, G. C. *et al.* Global DNA hypomethylation coupled to repressive chromatin domain formation and gene silencing in breast cancer. *Genome Res.* **22,** 246–258 (2012).
4. Song, Q. *et al.* A reference methylome database and analysis pipeline to facilitate integrative and comparative epigenomics. *PLoS One* **8,** (2013).
5. Heyn, H. *et al.* Whole-genome bisulfite DNA sequencing of a DNMT3B mutant patient. *Epigenetics* **7,** 542–50 (2012).
6. Sur, I. & Taipale, J. The NIH Roadmap Epigenomics Mapping Consortium. *Nat Rev. Cancer* **28,** 1045–1048 (2016).
7. Heyn, H. *et al.* Distinct DNA methylomes of newborns and centenarians. *Proc. Natl. Acad. Sci. U. S. A.* **109,** 10522–7 (2012).
8. Pacis, A. *et al.* Bacterial infection remodels the DNA methylation landscape of human dendritic cells. *Genome Res.* **25,** 1801–11 (2015).
9. Lund, K. *et al.* DNMT inhibitors reverse a specific signature of aberrant promoter DNA methylation and associated gene silencing in AML. *Genome Biol.* **15,** 406 (2014).
10. Li, Y. *et al.* The DNA Methylome of Human Peripheral Blood Mononuclear Cells. *PLoS Biol.* **8,** e1000533 (2010).
11. Newman, A. M. *et al.* Robust enumeration of cell subsets from tissue expression profiles. *Nat. Methods* **12,** 453–7 (2015).
12. Aran, D., Hu, Z. & Butte, A. J. xCell: Digitally portraying the tissue cellular heterogeneity landscape. *bioRxiv* 114165 (2017). doi:10.1101/114165
